# Nonlinear thinking in ecology and evolution: The case for ecological scaling of the Threshold Elemental Ratio

**DOI:** 10.1101/2024.08.20.608290

**Authors:** Benjamin B. Tumolo, Carly R. Olson, Erin I. Larson, Halvor M. Halvorson, Catherine E. Wagner, Amy C. Krist, Felicia S. Osburn, Eric K. Moody, Linnea A. Rock, Uchechukwu V. C. Ogbenna, Eli N. Wess, Briante Najev, Anthony J. Pignatelli, Jessica R. Corman

## Abstract

Nonlinear dynamics govern ecological processes, thus understanding thresholds is important for measuring and forecasting effects of climate change and management of natural resources. However, identifying whether and how such thresholds scale across biological levels of organization remains challenging. Ecological stoichiometry, the study of the balance of multiple elements and energy in ecological systems, provides a framework for scaling thresholds. We broaden a key organismal concept from ecological stoichiometry theory, the Threshold Elemental Ratio (TER), to study how nonlinear dynamics operate in evolutionary and ecological processes across the organizational hierarchy. Traditionally, TERs are used to describe the elemental ratio at which the limitation of organismal growth shifts from one element to another. Following this definition, we make a case for broadening the ecological scale of the TER beyond organisms to include populations, clades, communities, and ecosystems. We show how TERs can be detected and translated across different scales of biological and evolutionary organization through simulation modeling, literature review, and synthesis of empirical examples from diverse systems and ecological scales including: cyanotoxin production in lakes, alder-salmon dynamics, and the Cambrian explosion. Collectively, we demonstrate that TERs are widespread and consequential across levels of biological organization and that such thresholds manifest from a diversity of mechanisms. Thus, scaling of the TER concept holds promise for advancing our understanding of nonlinear dynamics from the micro-evolutionary to macro-ecological.

## Introduction

Thresholds in ecological and evolutionary relationships alter the trajectories of populations, communities, ecosystem processes, and evolutionary rates. Often the existence, location, and timing of thresholds is signaled by nonlinear relationships among ecological variables (Scheffer and Jeppesen 2007, Ratajczak et al. 2018, Spake et al. 2022). Thus, identifying the conditions required for thresholds to occur is foundational to understanding dynamic ecological processes and systems. For example, understanding thresholds is critical to measuring and forecasting the effects of climate change and for management of natural resources (Dodds et al. 2010, deYoung et al. 2008). Despite significant progress in identifying the causes and consequences of thresholds to ecological and evolutionary processes, the challenge of how to unify these phenomena across biological levels of organization remains.

Ecological stoichiometry—the study of the balance of multiple chemical elements and energy in ecological systems — is an attractive framework for translating nonlinear dynamics and shifts in nutrient limitation across scales because it characterizes interactions between organisms and their environment using a common currency (i.e., chemical elements and energy) grounded in the first principle of conservation of mass and energy (Sterner & Elser 2002). A key concept from ecological stoichiometry, the Threshold Elemental Ratio (hereafter TER, Olsen et al. 1986; Urabe and Watanabe 1992) describes the nonlinear change in organismal growth or fitness when resource limitation switches from one element to another (Sterner and Elser 2002; Frost et al. 2006). The TER concept is rooted in stoichiometric models that couple organismal bioenergetics and body elemental composition (Frost et al. 2006). Yet, despite few examples where the TER has been considered in communities (e.g., Moody et al. 2019, Elser et al. 1998) and ecosystem properties (e.g., Mooshammer et al. 2014, Elser and Urabe 1999), TERs have predominantly been applied at the organismal scale. However, because all organisms require carbon (C), nitrogen (N), phosphorus (P), and other elements for growth and metabolism, insufficient supply of one or more of these essential elements is expected to control process rates from biogeochemical transformations to community structure (Urabe & Watanabe 1992; Elser et al. 2007, Schade et al. 2011, Tromboni et al. 2018), as well as rates of microevolutionary and macroevolutionary processes (Elser et al. 2006, Frisch et al. 2014). Despite the potential for nonlinear shifts in nutrient limitation to occur across ecological scales, very few works have examined this possibility via ecological stoichiometry theory, or specifically TERs. Given that the TER concept couples nonlinear change and the effects of relative amounts of elements on biological processes, we suggest that expanding the TER concept beyond the organismal scale will provide testable predictions that could advance our understanding of nonlinear dynamics driven by shifts in nutrient limitation across ecology and evolution more broadly.

Here, we broaden the TER concept across scales and show how nonlinear stoichiometric controls strongly influence ecological processes across biological levels of organization. We assert that broadening the TER concept is needed to integrate the fundamental contributions of ecological stoichiometry across evolutionary and ecological processes at multiple levels of organization. To make this case we begin by providing a background on the application of TERs to date. Then, we demonstrate the prevalence and importance of TERs from a diversity of ecosystems and ecological and evolutionary processes by compiling and describing case studies from the literature. Next, we present results from a literature review and a simulation model aimed at evaluating the applicability of scaling the TER concept to contemporary ecological questions across biological levels of organization. We conclude by providing recommendations and considerations for broadening the TER concept and highlighting areas of future research.

### Ecological scaling of the Threshold Elemental Ratio

Our scaling of the TER concept is based on the ratio of elements at which an organism, population, community, and/or ecosystem exhibits a nonlinear change driven by a shift in limitation in an ecological or evolutionary state or process (Figure 1). Thus, our definition of a TER moves beyond organisms to be applied to and include populations, clades, communities, ecosystems, and macrosystems. As such, the TER should pertain to a wide breadth of time frames (physiological to evolutionary) and spatial scales (microscopic to global) (Table 1, Table S1). For example, a TER may reflect a shift in the element most limiting to community level coexistence among competitors (Tilman 1982) or ecosystem processes (Mooshammer et al. 2014). Ecological scaling of the TER provides a means of explaining nonlinear responses in diverse situations where the currencies of available matter and energy drive the response. Mathematically, the TER constitutes the ratio at which an inflection point, threshold, or break occurs in a relationship between the stoichiometric driver and response variable (Figure 1a). Thus, identifying and quantifying TERs at various scales can be straightforward and rigorous. As evidenced by many studies, nonlinear ecological dynamics are common (Dodds et al. 2010; Spake et al. 2022), and many of these relationships, when they involve changes in stoichiometric ratios, may qualify as a TER, but are not captured within the scope of the current framework (Table 1, Table S1). Integrating these existing examples with a broader TER concept should improve our understanding of nonlinearity in ecology and evolution in a synthetic and scale-inclusive manner with shifts in nutrient limitation as the common underlying mechanism.

**Fig 1.**
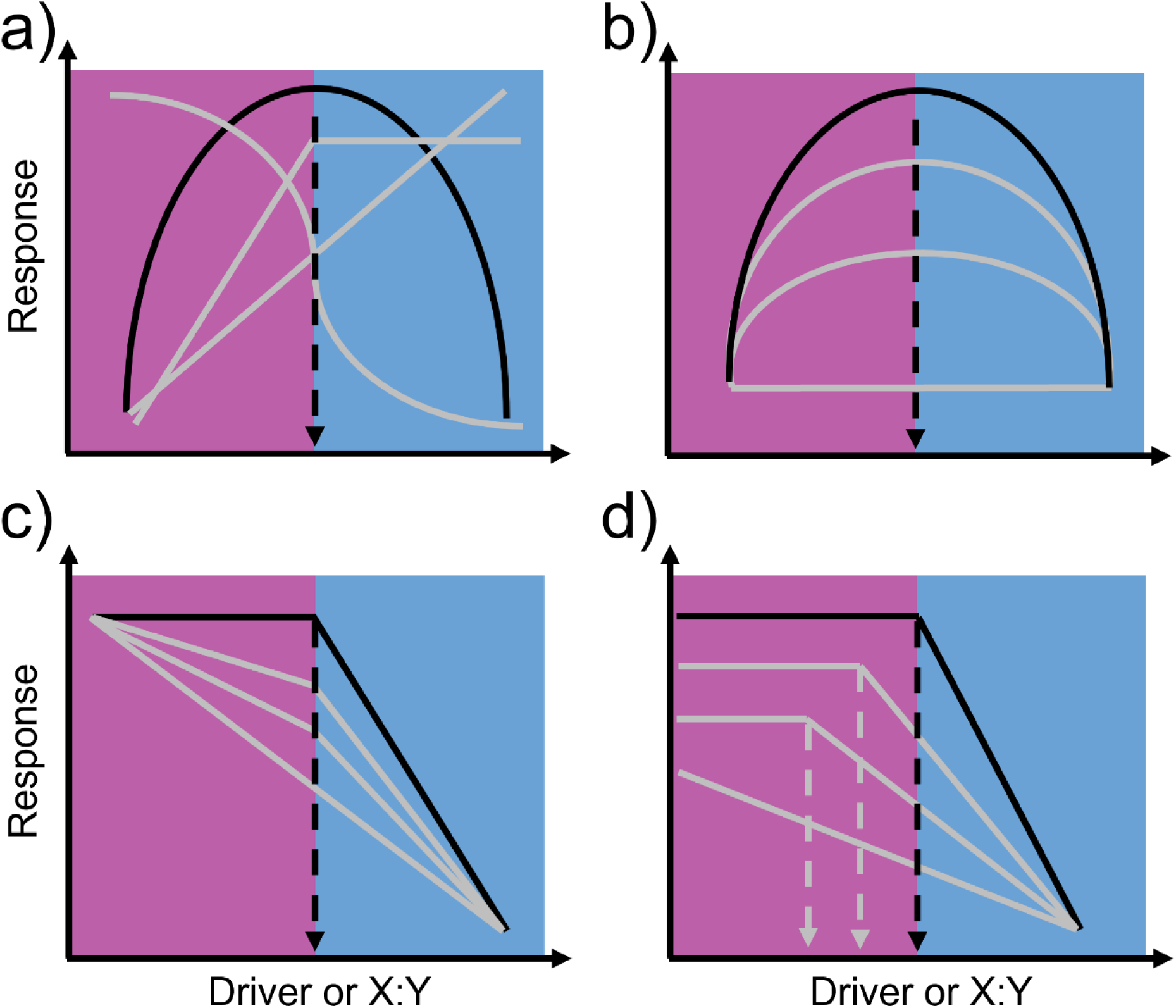
Conceptual visualization of scaling the threshold elemental ratio (TER). Focal TER relationships are black, and theoretical variations from the focal TER are gray. Shading in panels indicates ranges below (purple; left) or above (blue; right) the focal TER. a) The TER may result from a diversity of potential nonlinear relationships between the driver or X:Y ratios (indicating the ratio of two elements X and Y) and the response; b) TER scenarios may differ in overall sensitivity above and below the TER; c) Across different scenarios, the degree of linearity vs. nonlinearity may also differ; and d) the TER itself may differ, depending on system attributes or environmental drivers.

**Table 1:** Examples of elemental stoichiometry driving nonlinear thresholds at various ecological, evolutionary, spatial, and temporal scales. Grain Temporal is the temporal scale of individual observation units; Grain Spatial is the spatial scale of individual observation units; Extent Temporal is the temporal scale of the entire study; Extent Spatial is the spatial scale of the entire study. Abbreviations: “Level of Org.”, Level of biological organization (organismal, community, ecosystem); “P”, predictor; “R”, response; “stoich”, stoichiometry.

| Topic | Elements | Ecosystem | Driver | Level of Org. | Grain |  | Extent |  | Response | Reference |
| --- | --- | --- | --- | --- | --- | --- | --- | --- | --- | --- |
|  |  |  |  |  | Temporal | Spatial | Temporal | Spatial |  |  |
| Adaptive radiation of *Geissois* sp. | Ni, Al, P, Mg, Ca, Co, Cr, Fe. | Terrestrial: tropical forest | Ultramafic soil | **P:** Ecosystem (soil)<br>**R:** Community (speciation rate) | NA | Species within a genus | 7 m. years | New Caledonia | Threshold | Pillon et al. 2014 |
| Global leaf decomposition | C, N, P | Aquatic and terrestrial | Vegetation stoich | **P:** Organismal (Species C:N:P)<br>**R:** Ecosystem (decomposition rate) | NA | NA | Global | Global | Loglinear | Enriquez et al. 1993 |
| Streamwater chemistry affect periphyton community | C, N, P | Aquatic: experimental stream | Streamwater stoich | **P:** Ecosystem (Water N:P ratio)<br>**R:** Ecosystem (periphyton N:P, C:N, C:P) and Community (chlorophyll) | 3 sampling days | Clay tile (0.5 mm<sup>2</sup>) | 28 days | 30 flumes (2.8mx10cm) | Threshold | Stelzer & Lamberti, 2001 |
| Microbial efficiency | C, N | Terrestrial: soils | Soil N availability | **P:** Ecosystem (resource, C:N, stoich)<br>**R:** Population (N use efficiency) | one time point | Within same latitude and longitude | 5 years | Global | Threshold | Mooshammer et al. 2014 |
| Invasive fish effect river water N and P | N, P | Aquatic: rivers | Invasive fish excreting at high N:P | **P:** Organismal/Population (abundance)<br>**R:** Ecosystem (N and P availability) | hrs-days | Grab samples | hrs-yrs | 550m reach | Threshold | Capps and Flecker 2013 |
|  |  |  |  |  | Temporal | Spatial | Temporal | Spatial |  |  |
| Cambrian explosion | P | Aquatic: Proterozoic - Cambrian oceans | Phosphogenic event | **P:** Ecosystem (C:P)<br>**R:** Organismal (growth rates, metabolism, evolution of skeletons) | NA | NA | 20-30 m. years | Global | Threshold | Cook 1992 |
| Database modeling of CNP influence on soil bacterial diversity | C, N, P | Terrestrial (soil) | Resource organic soil matter C:N:P | **P:** Ecosystem (soil C:N:P ratio)<br>**R:** Community (Shannon Diversity) | Single time point | 20x20 km<sup>2</sup> sampling grid | 4 yrs | 6 ecosystem types | Threshold | Delgado-Baquerizo et al. 2017 |
| Organic matter stoich and stream N, P uptake | N, P | Aquatic: forested stream | Organic matter N:P | **P:** Ecosystem (organic matter stoichiometry)<br>**R:** Ecosystem (N:P uptake) | Single time point | 10-20 m | 1 yr | 160m | Threshold | Gibson and O'Reilly 2012 |
| Soil microbial carbon-use efficiency | C, N | Aquatic: soils | Organic matter N | **P:** Ecosystem (C:N nutrient availability)<br>**R:** Ecosystem (carbon-use efficiency) | NA | NA | NA | Global | Threshold | Manzoni et al. 2012 |
| Eutrophication destabilizes ecosystem | N, P | Aquatic: lakes | Increased P availability | **P:** Ecosystem (amount of nutrients in the sediment)<br>**R:** Community (shift in macrophyte) | Single time point | 0.2m<sup>2</sup> | 2 years | 46 lakes in the Yangtze river basin, China | Threshold | Su et al 2019. |
| Ecosystem N processing rates | C, N | Ocean, great lakes, and streams | Increased resource organic carbon | **P:** Ecosystem (organic carbon:nitrate ratio in water)<br>**R:** Community (microbial production) | NA | NA | Multiple years (data from multiple databases) | Global | Threshold | Taylor and Townsend 2010 |

Many factors are expected to determine the occurrence and characteristics of stoichiometrically driven thresholds at various scales of inquiry. Some systems, may exhibit more sensitivity to thresholds driven by stoichiometry compared to others (Figure 1b): for example, in a synthesis of soil microbial carbon use efficiencies, sensitivity to substrate C:N was lower in low-fertility systems than high-fertility systems, likely because of an overall limitation of maximum carbon use efficiency under low fertility (Manzoni et al. 2012). Within a single study system, a threshold may occur at one biological scale but not at another; this mismatch may reflect differing timeframes required for the threshold to take effect, or it may reflect decoupling among variables. As an example, in an experiment using the herbivorous zooplankter *Daphnia magna* fed a wide dietary C:P gradient, *Daphnia* sp. growth exhibited a nonlinear threshold response of growth rates that peaked at intermediate C:P; yet, feeding rates decreased linearly with decreasing diet C:P (Plath and Boersma 2001). Likewise, evolutionary responses may be linear at the scale of genotypic response, but nonlinear at the level of clade-level diversification rate (Figure 2). In this way, the same driver may elicit a nonlinear effect on one response variable but a linear effect on another or, the nature of the nonlinearity (e.g., threshold, degree of nonlinearity) may differ between two responses to the same driver (Figure 1a & c). Finally, the threshold itself may depend on system attributes or environmental drivers (Figure 1d) as evidenced by shifts of organism TERs in response to temperature (Laspoumaderes et al. 2022). Ultimately, parsing the nature and determinants of stoichiometrically-driven thresholds, or shifts in limitation, across scales presents fertile theoretical ground for further study.

**Fig 2.**
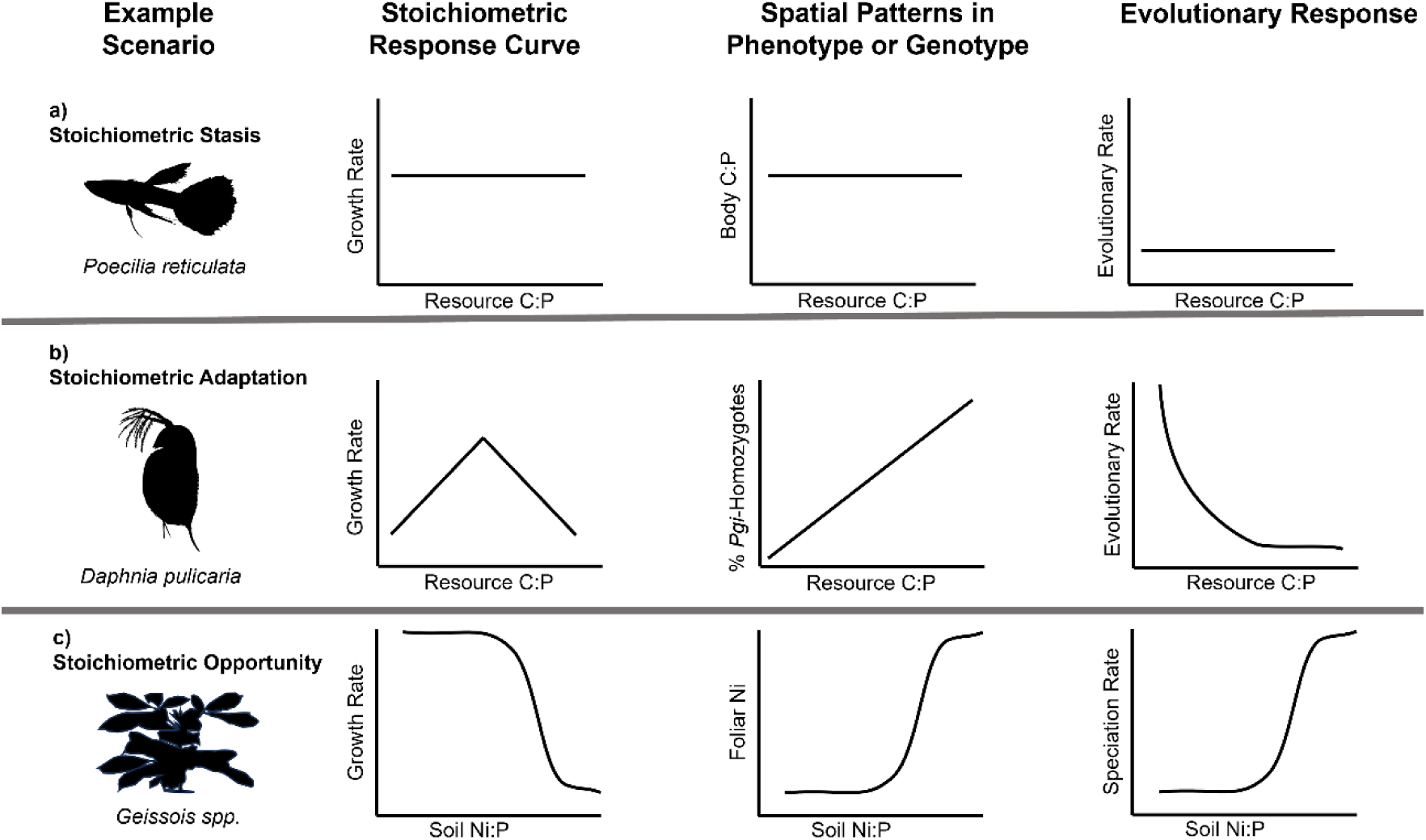
Example response curves for resource stoichiometry and the adaptive responses that might result from them. Plots depict fitness response curves of individual organisms, patterns in phenotype or genotype among populations at a given time, and patterns in evolutionary rate for a population as a function of stoichiometric drivers. a) Fitness, phenotypic/genotypic, and evolutionary rate response curves are all invariant with resource stoichiometry; there is no threshold on evolutionary processes. b) Fitness exhibits a unimodal or knife-edge relationship with resource stoichiometry. Phenotypic or genotypic responses vary linearly over space, but evolutionary rate varies over time; thus, a threshold exists for organismal performance and for evolutionary rate, but is not apparent when sampling over space. c) Nonlinear threshold fitness response curves drive nonlinear responses for the phenotype or genotype and for evolutionary rates. A threshold is evident across temporal and spatial scales.

### Case studies

We compiled and described several case studies to demonstrate the transferability of TERs to a diversity of ecosystems and ecological and evolutionary processes (Table 1, Table S1). This compilation includes organismal examples ranging from microbes to trees and ecosystem functions from N-cycling to organic matter processing, in both terrestrial and aquatic habitats. Below, we expound on three examples from Table 1 to illustrate the TER concept at different scales and describe how TERs can influence micro-and-macroevolutionary processes (Figure 2).

### Lake eutrophication and cyanotoxin production

Absolute concentrations of N and P have long been recognized for their role in lake eutrophication. Importantly, these dynamics may represent ecological scaling of the TER concept as the relative availability of N and P often mediate shifts in lake productivity and eutrophication. For instance, constraints on N supply likely limit phytoplankton growth in P-replete Canadian prairie lakes (Hayes et al. 2019). In contrast, increased diffuse watershed-based N inputs likely increased productivity in P-rich lakes in Northern Ireland (Bunting et al. 2007). Lakes in watersheds with greater human activity may accumulate P faster than N (Yan et al. 2016), leading eutrophic lakes to have a lower total N:P ratio than less nutrient-rich ecosystems (Zhou et al. 2022).

These shifts in lake N:P stoichiometry may drive nonlinear dynamics in the production of harmful cyanotoxins and thus represent an ecosystem scale TER with far-reaching management implications. Cyanotoxins, secondary metabolites produced by cyanobacteria, can affect water quality and threaten human and other organism health (Buratti et al. 2017, Lopez-Rodas et al. 2008). The chemical structure of cyanotoxins can vary from N-rich to C-rich, with molar C:N ranging from about 2-43 (Van de Waal et al. 2014). This variation in cyanotoxin structure sets up the potential for a TER. Specifically, the cyanobacteria *Microcystis* spp. produces more cyanotoxin microcystin under lower water column C:N (Van de Waal et al. 2009, Osburn et al. 2022) and microcystin production can decrease non-linearly with *Microcystis* spp C:N and C:P tissue stoichiometry, suggesting a shift from C- to N or P limitation may drive nonlinear cyanotoxin production. Other opportunities to scale TERs of cyanotoxin production exist beyond *Microcystis* spp. and freshwaters. For example, *Aphanizomenon* spp. (Wagner et al. 2023) and *Dolichospermum* spp. (Kramer et al. 2022) produce more cyanotoxins under N-rich environmental conditions. In contrast, C-rich cyanotoxins (e.g., karlotoxins and amnesic shellfish poisoning toxin) found in oceans generally increase under conditions of N- or P-limitation (Van de Waal et al. 2014). These observations prompt multiple questions: do these genera demonstrate thresholds with respect to cyanotoxin production and relative nutrient availability? If so, what are the thresholds and how do they compare to *Microcystis* spp.? How does the chemical composition (i.e., N-rich vs. C-rich) of the cyanotoxin influence the shape and location of the threshold? Importantly, detecting a TER of cyanotoxin production based on lake N:P concentrations may be hampered by the N-fixing capabilities of many cyanobacteria which increase cellular N concentrations independent of environmental N supply (Osburn et al. 2022). Nevertheless, understanding the relative availability of N and P has been instructive to predicting and managing harmful algal blooms. Thus, placing this understanding within a TER may provide path forward by focusing on drivers of shifts in limitation of cyanotoxin production, whether these drivers are acting on environmental supply or species demand for energy and nutrients.

### Alder and salmon influences on nitrogen dynamics

In terrestrial ecosystems, plants that host N-fixing symbionts can play a large role in the relative availability of N to C or P (Vitousek et al. 2002). Alder (*Alnus* spp.) are shrubs that form a symbiotic relationship with N-fixing bacteria. Because this symbiosis results in abundant inorganic N in the soil near extensive patches of alder, streams in catchments with high percent cover of alder experience large amounts of N export from their surrounding watershed (Stottlemeyer and Toczydlowski 1999). Particularly in boreal and coastal rainforest headwater streams that lack marine-derived N from migratory salmon (Shaftel et al. 2012), alder may promote P limitation of aquatic ecosystem processes through N-saturation (Shaftel et al. 2012, Devotta et al. 2021), although nutrient limitation by N and P may also depend on algal community composition (Volk et al. 2008). Alder cover is also related to N fixation rates, with in-stream dissolved inorganic N linearly increasing with alder cover in the catchment and benthic N fixation rates linearly decreasing with increasing alder cover (Hiatt et al. 2017).

While increases in alder cover are associated with linear trends in some ecosystem responses among streams, shifts in alder cover and their resultant effects on stream ecosystem stoichiometry can create thresholds by generating nonlinear dynamics over time. Particularly under a warming climate, alder expansion could result in higher levels of N export from watersheds (Salmon et al. 2019), where the amount of alder cover in the watershed initiates a threshold response by increasing stream water N:P. This nonlinear response of stream water N:P to alder cover may, in turn, influence microbial metabolic activity, via a shift from N- to P-limitation (Devotta et al. 2021). Alder cover can also initiate stoichiometric-driven thresholds through interactions with other terrestrial plant species. When the effects of salmon marine-derived N on white spruce were investigated at sites with and without alder, Helfield & Naiman (2002) found that foliar C:N in white spruce (*Picea glauca*) was highest at sites without alder or salmon present, and alder and salmon interacted to influence white spruce basal area growth.

### The Cambrian explosion

The end of the Proterozoic eon (∼ 540 MYA) was marked by extensive erosion of terrestrial material into the ocean, causing influxes of calcium (Brennan et al. 2004), phosphate (Cook 1992, Brasier and Callow 2007), and other ions (iron, strontium, bicarbonate) into the oceans that exceeded any previous period of Earth’s history in the geological record (Squire et al. 2006). These nutrients triggered an explosion of algae and cyanobacteria (Squire et al. 2006) which produced a large increase in oxygen from photosynthesis (Campbell and Squire 2010). The combined effects of increased oxygen, calcium, and phosphate causing a shift in, or release from, elemental limitations may have been a driving mechanism for the Cambrian faunal “explosion” and the initiation of biomineralization (Elser et al. 2006). This explosion resulted in a threshold response of the largest diversification event of animals at the global scale.

Dramatic changes in calcium and P availability facilitated major transitions in the evolution of animals, the evolution of exoskeletons, and possibly the evolution of higher growth and metabolic rates of metazoans (Elser et al. 2006) and the evolution of self, non-self recognition (Fernàndez-Busquets 2010). Increased phosphate and calcium levels may have favored the evolution of phosphatized and calcified exoskeletons. Hydroxyapatite (Ca_10_(PO_4_)_6_(OH)_2_), the primary constituent of vertebrate bones, detoxifies excess levels of P and its expanded use as a structural material may have facilitated increased diversification rates during this era (Cohen et al. 2017). Similarly, selection to reduce toxicity of calcium ions to normal cell function may have caused the evolution of existing biochemical pathways to mineralize calcium. Shells composed of calcium carbonate (CaCO_3_) detoxify both calcium and oxygen ions, and supersaturation of CaCO_3_ permitted animals in all phyla to precipitate CaCO_3_ for the first time (Squire et al. 2006) consistent with the first appearance of skeletal marine phyla from ∼545 Ma (Martin et al. 2000). At the beginning of the Cambrian period, large increases in P, the most limited nutrient in the oceans over geological timescales, in concert with increased oxygen levels, would have permitted higher animal growth rates by relieving severe P limitation (Elser et al. 2006). Calcium increases the binding forces between calcium-dependent cell adhesion molecules, thus large increases in calcium concentrations could have contributed to the Cambrian explosion by increasing adhesion between cells, thus facilitating the evolution of self, non-self recognition, and possibly multicellularity (Fernàndez-Busquets 2010).

Vast scale geological processes occurring at the end of the Phanerozoic permitted cascading and nested geochemical and biological feedback loops (e.g., evolutionary arms race, evolution of Hox genes) that together generated the Cambrian explosion (Smith and Harper 2013). Thus, stoichiometric constraints alone cannot explain this diversification event, but geological processes that released elements initiated the chain of events that led to the Cambrian explosion (Smith and Harper 2013), and elemental availability had fundamental impacts on the directions of further diversification. The stoichiometric drivers of the Cambrian explosion were unprecedented global increases in oxygen, calcium, and P, leading to the largest diversification event of animals. By any measure, the Cambrian explosion represents a nonlinear increase in diversification rate (Figure 2c) influenced by stoichiometric shifts in resource limitation; this evolutionary event had fundamental lasting impacts on the earth’s biodiversity.

### Evolutionary thresholds

TERs may exist in an evolutionary context when organismal fitness or diversification vary with the stoichiometry of resources or the environment in a way that facilitates nonlinear evolutionary responses. At a microevolutionary level, components of fitness, such as somatic growth rate and fecundity, can vary with resource stoichiometry via the TER. The shape of growth or fecundity reaction norms over gradients in resource or environmental stoichiometry can therefore dictate whether a threshold on evolutionary responses will exist. For example, if an organism’s fitness does not vary with resource stoichiometry, it is unlikely that an evolutionary TER acts on that organism (Figure 2a). Trinidadian guppies, for instance, vary in body stoichiometry among populations, but diet stoichiometry does not appear to have a large influence on growth rates or body stoichiometry (El-Sabaawi et al. 2012; Figure 2a).

If fitness is optimized at a single stoichiometric ratio with symmetric decreases in fitness away from this optimum TER, phenotypic or genotypic responses may be linear when sampled over a spatial gradient in stoichiometry (Figure 2b). However, even if spatial patterns suggest a lack of a threshold, evolutionary rate may still vary with resource stoichiometry in a nonlinear fashion (Figure 2b). One example where this may occur is in the zooplankton *Daphnia pulicaria*, whose growth and fecundity are limited at both high and low C:P. Growth rates and fecundity in populations of *Daphnia* may vary in their TER over both time and space due to agricultural intensification promoting low resource C:P; however, the percentage of the population comprised of homozygotes at a gene linked to P use efficiency, *Pgi*, to resource C:P appear to be linear (Moody et al. 2022; Figure 2b). Nonetheless, resurrection of resting eggs from dated sediment cores has revealed nonlinear dynamics in the rate of evolutionary change over time within the population of *Daphnia pulicaria* from a lake that became increasingly eutrophic over time (Frisch et al. 2014; Figure 2b).

Asymmetric reaction norms could lead to both nonlinear phenotypic responses over spatial gradients and evolutionary responses over time (Figure 2c). Trees in the genus *Geissois*, which colonized the island of New Caledonia, radiated into thirteen species from a single common ancestor. In New Caledonia, *Geissois* spp. encountered ultramafic soils with relatively high amounts of toxic metals like nickel (Ni) and low amounts of nutrients like N, P, and K. Above the threshold for Ni toxicity, *Geissois* spp. populations evolved several distinct strategies for surviving on these soils, ultimately leading to an increase in speciation rate in these environments (Pillon et al. 2014; Figure 2c). However, it is not possible to disentangle whether this nonlinear evolutionary response was solely driven by stoichiometry per se or the ecological opportunity associated with colonizing novel environmental characteristics. Links between micro- and macroevolutionary processes can be complex and discontinuous (Gould 1998, Simons 2002). However, if influences on organismal fitness also have implications for speciation or extinction rates, they can influence macroevolutionary patterns (Rabosky and McCune 2010). The logic of the *Geissois* tree example follows this pattern – local adaptation to diverse stoichiometric conditions could accelerate speciation rates as well as influence adaptation within populations. Direct empirical evidence of stoichiometric shifts in resource limitation as a mechanism for evolutionary thresholds remains quite limited and represents a promising direction for future research.

### Literature review

We conducted a systematic literature review to evaluate the utility of scaling the TER. Our literature review identified the extent to which ecological papers that focused on two essential elements, N and P, also considered the stoichiometry of these elements. We gathered papers using the search terms “nitrogen AND phosphorus” from the journals “Ecology Letters”, “Ecology”, “Functional Ecology”, and “Oikos” from the Web of Science in December 2022. We used these journals as a representative, albeit non-exhaustive, sampling of the ecological literature. The search generated 663 papers and included papers published from the years 1954-2022. We examined and read the titles and abstracts to assess suitability (i.e., “was the paper an empirical study that focused on nutrients?”) with 563 considered appropriate. We then examined suitable studies more closely and answered the following questions: (1) Did the study measure a nutrient (e.g., N or P)? (2) Did the study measure both N and P? (3) Did the study report a stoichiometric ratio? (4) Was a stoichiometric ratio used in a statistical test? (5) Was a stoichiometric ratio used as (a) an explanatory variable, (b) a response variable, or (c) both?

We found that all studies included a topic relevant to nutrients in ecological systems. A small number of studies (n=63, 11%) only examined a single nutrient, and therefore were unable to test stoichiometric questions (Figure 3). A greater proportion of papers measured multiple elements, but did not present or analyze these elements as stoichiometric ratios (n=237, 42%). Therefore, the studies within this category missed an opportunity to explore TERs despite possessing the necessary information to do so (Figure 3). Nearly half of the studies (n=263, 47%) explored TERs or could have explored extended TERs by using a stoichiometric ratio as an explanatory and/or response variable (Figure 3). Although our consideration of the TER is focused on the stoichiometric ratio as the explanatory variable, we included both explanatory and response variables in this analysis as a more comprehensive estimation of studies that had the potential to investigate the concept. Overall, this finding indicates that the TER concept can unify and be meaningfully applied to a considerable portion of existing nutrient-focused ecological literature and possibly catalyze future research on the topic. The database for the literature review can be accessed here: https://doi.org/10.5281/zenodo.12628550.

**Fig 3.**
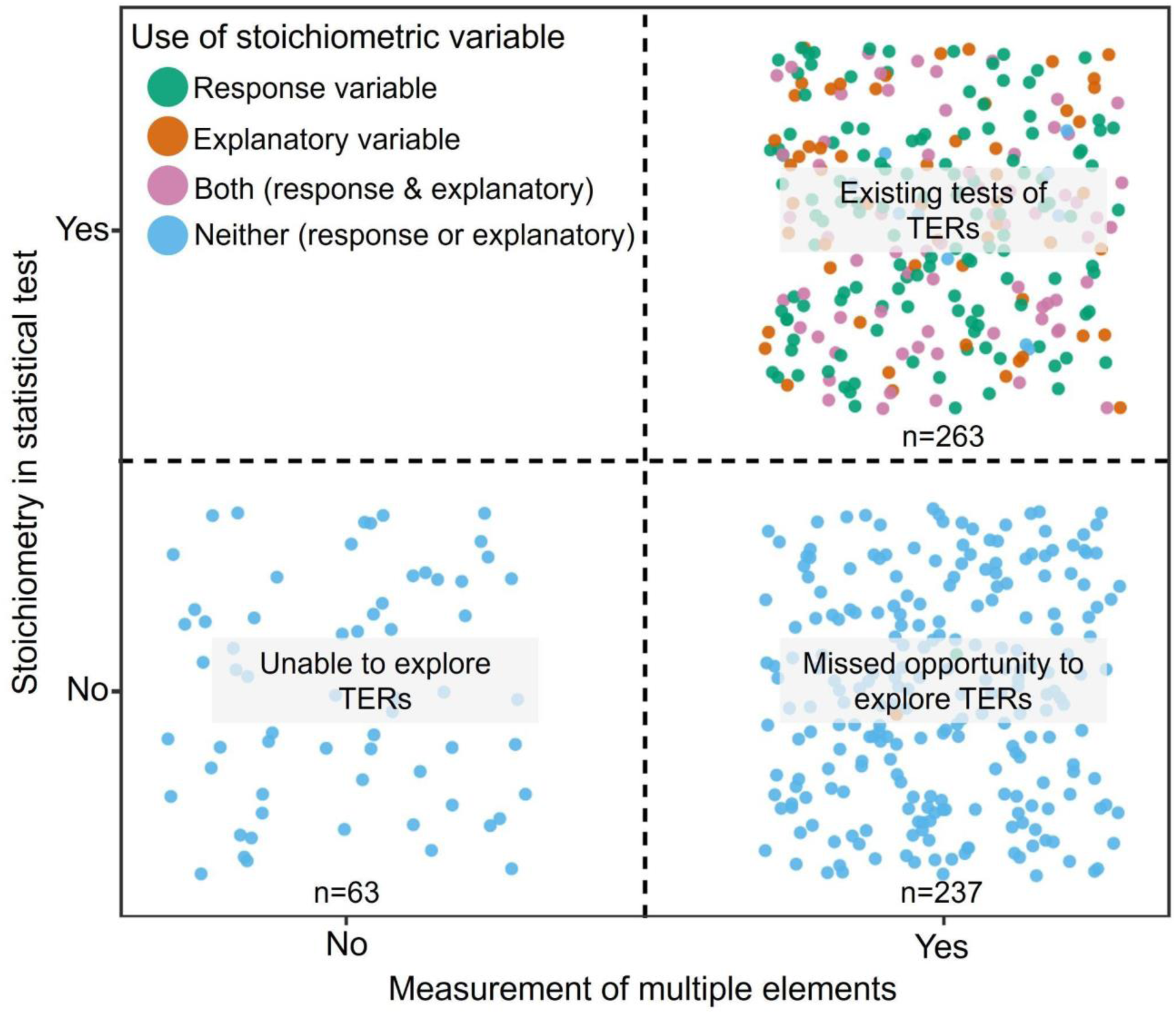
Ecological studies focused on nutrients and their potential to address threshold elemental ratio (TER) scaling from the literature. Studies are plotted based on whether multiple elements were measured and if elements were examined statistically as a ratio. Color refers to the use of variable within models. Text overlay provides interpretation of studies relevance to the TER concept based on placement within the plot.

### Modeling Scaled Threshold Elemental Ratios

Here we provide an example analysis of an ecosystem scale TER using a lake ecosystem process model embedded with an algal physiological model. Using this process model, we evaluated the response of lake ecosystem algal biomass to a gradient of supply N:P stoichiometry (See Figure 4 for methodological details). Our goals with this model were to: 1) demonstrate the presence of a TER for an ecosystem-scale response using algal biomass as an indicator of ecosystem autochthonous POC pools; and 2) provide examples of mechanisms acting across scales that may drive variation in the shift in limitation underlying the TER. Specifically, we were interested in how the location and response type (i.e.; x- and y-value) of the threshold may vary under different parameterizations using a well-established theory (Figure 1; Klausmeier et al. 2008).

**Fig 4.**
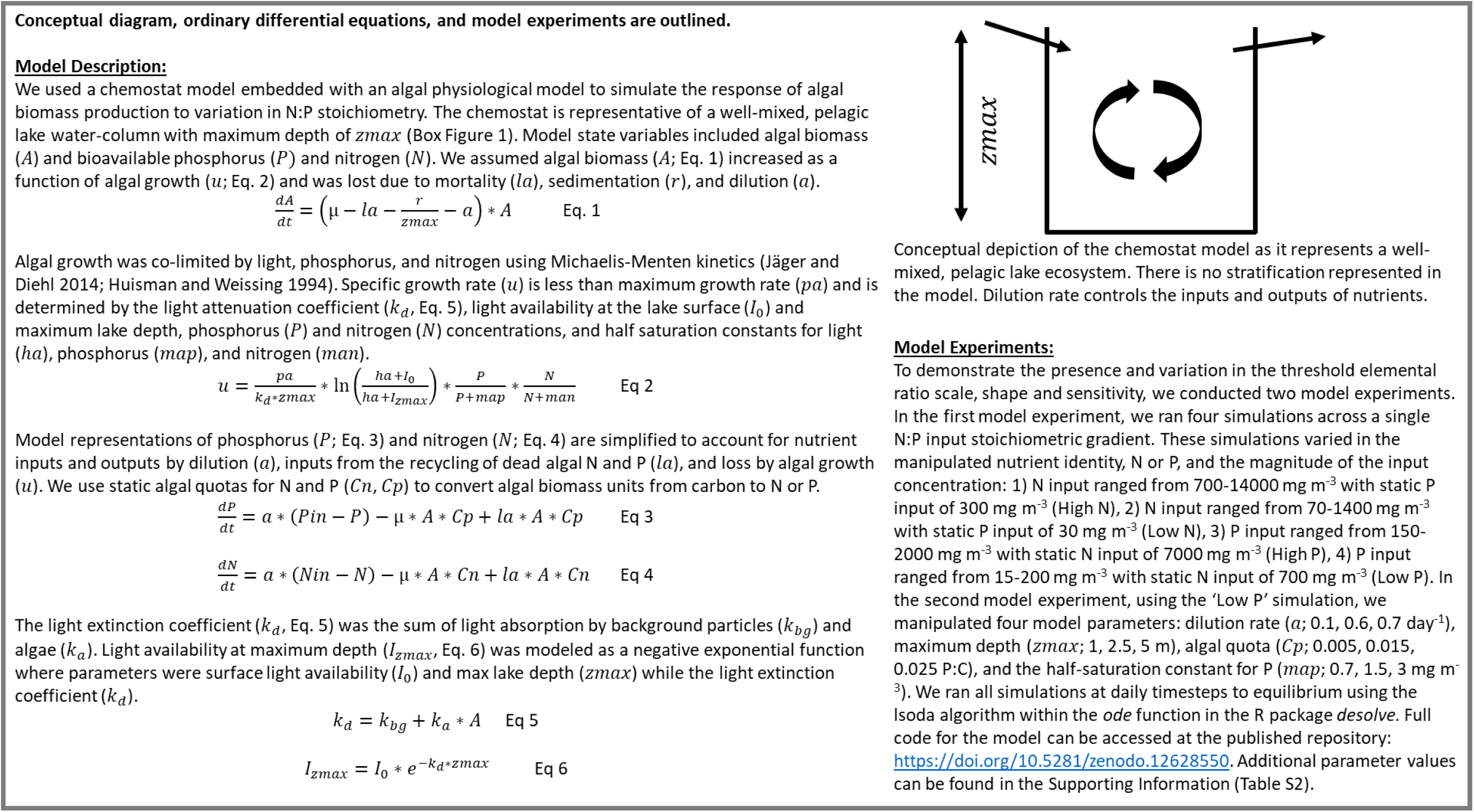
Conceptual diagram, ordinary differential equations and outline of modeling experiments.

Model simulations revealed that algal biomass responded non-linearly to a gradient of supply N:P, indicating the presence of a TER driven by shifts in limitation between N and P (Figure 5, Supplementary Figure S1). The shape of the threshold response type was influenced by whether N or P was manipulated, and sometimes the absolute concentration of nutrients (Figure 5). When N was manipulated, the response type of the TER was described by a saturating threshold independent of the absolute concentration caused by a shift from N to P limitation (Figure 5a & 5b, Supplementary Figure S1). Thus, when the numerator was manipulated, the response type was robust to variation in absolute concentration. In contrast, manipulating high absolute concentrations of P resulted in the absence of a TER despite the presence of a shift from N to P limitation (Figure 5c, Supplementary Figure S1). The absence of a TER is due to high P concentrations switching the primary limitation on algal biomass accrual from nutrients to light via algal self-shading. In the low P manipulation, the response type was once again a threshold across a stoichiometric gradient (Figure 5d). The stoichiometric location (x-value) of the threshold was agnostic to both the identity and magnitude of the nutrient supplied and was always ∼16 N:P. However, the response threshold (y-value) did respond to nutrient identity and magnitude (Figure 5). These results are consistent with previous research (Downing and McCauley 1992, Bergstrom 2010); a stoichiometric ratio of 16 is customarily used as the threshold between N and P limitation for algal dynamics (Redfield 1958). However, system-specific nuances, such as community composition, often lead to deviations from this broader generalization of 16 and model parameterization of algal stoichiometric traits can be implemented to reflect this (Figure 6; Reynolds 1992, Klausmeier et al. 2004).

**Fig 5.**
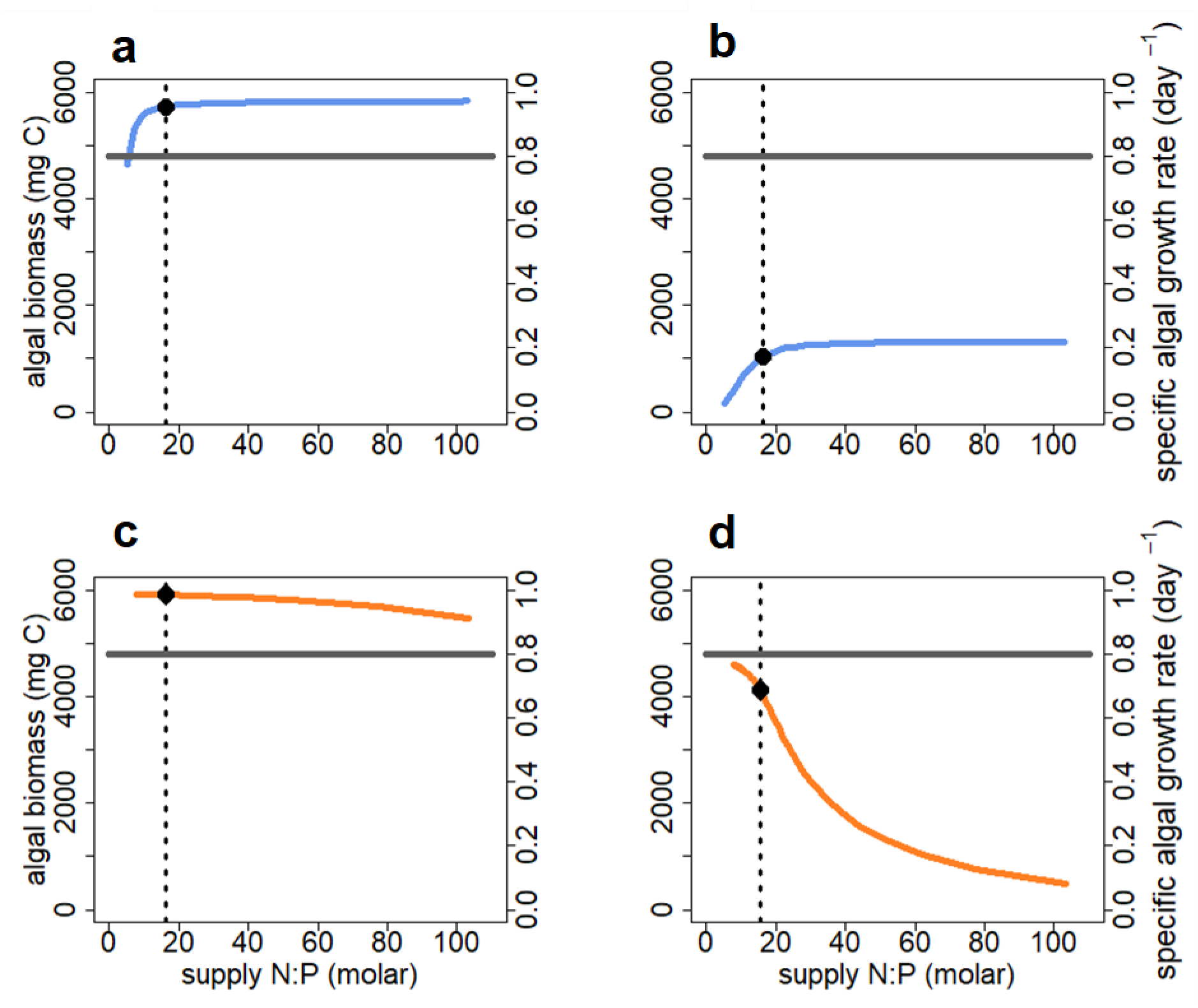
Results from the first model experiment to demonstrate how the identity and magnitude of the manipulated nutrient (N vs P) alters the shape, sensitivity, scale, and location of the threshold elemental ratio (TER) describing the response of lake algal biomass (mg C) to a supply N:P stoichiometric gradient. a) N input ranged from 700-14000 mg m^-3^ with static P input of 300 mg m^-3^ (High N, solid blue line), b) N input ranged from 70-1400 mg m^-3^ with static P input of 30 mg m^-3^ (Low N, solid blue line), c) P input ranged from 150-2000 mg m^-3^ with static N input of 7000 mg m^-3^ (High P, solid orange line), d) P input ranged from 15-200 mg m^-3^ with static N input of 700 mg m^-3^ (Low P, solid orange line). The dotted line in all panels marks the stoichiometric location (x-value) of the threshold. The solid gray line marks algal specific growth rate (day^-1^). A TER occurred at the ecosystem scale (algal biomass response), but not at the organismal scale (growth rate response). The solid point is the x- and y-value of the TERs.

**Fig 6.**
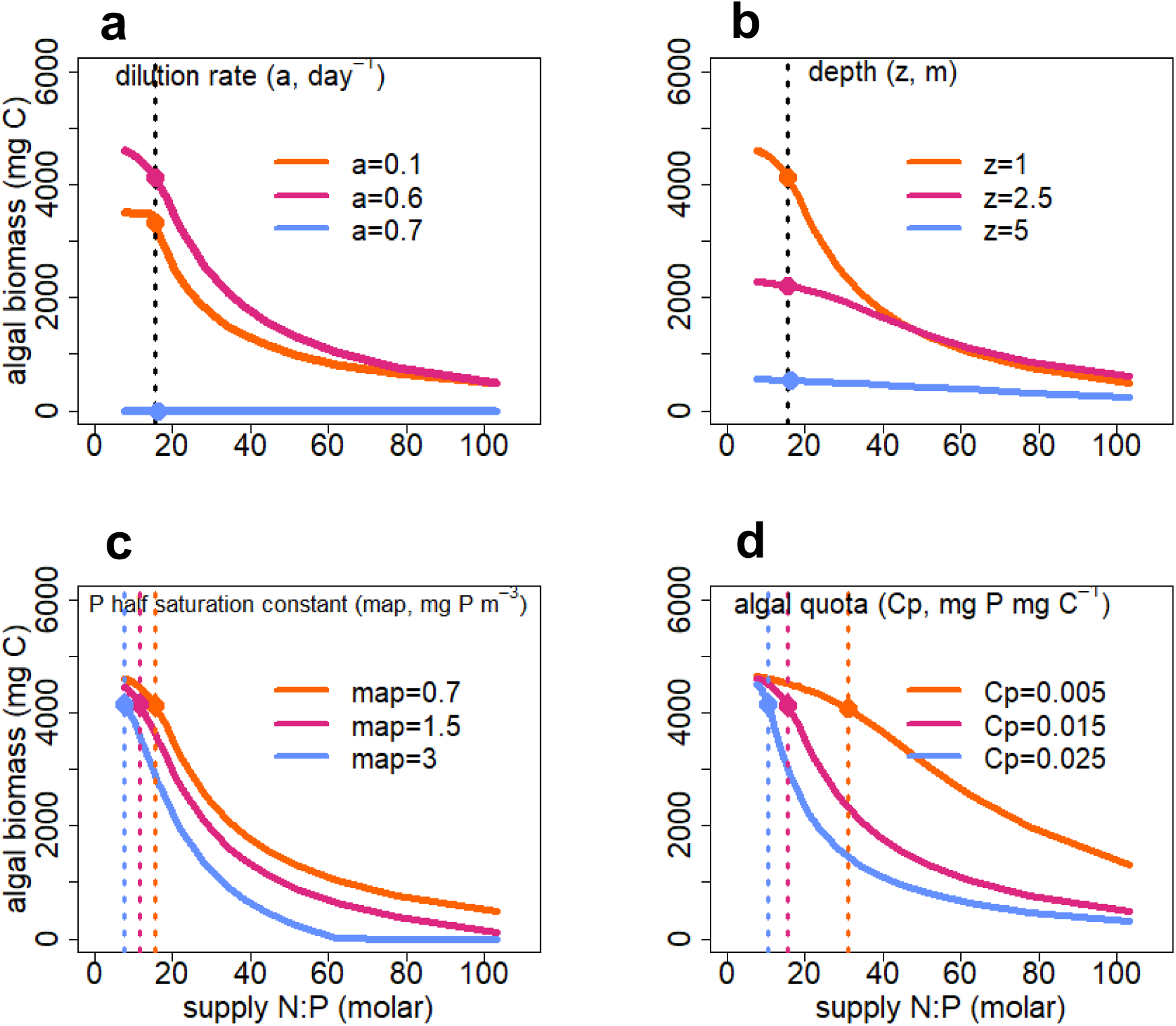
Using the “low P” simulation from the first model experiment, results from the second model experiment demonstrated how different mechanisms acting across scale can alter the shape, and shift the stoichiometric location (x-value) and y-value, of the TER. a) dilution rate (a; 0.1, 0.6, 0.7 day^-1^), b) maximum depth (zmax; 1, 2.5, 5 m), c) algal quota (Cp; 0.005, 0.015, 0.025 P:C), d) the half-saturation constant for P (map; 0.7, 1.5, 3 mg m^-3^). The dotted lines mark the stoichiometric location of the TER. The solid points are the x- and y-values of the TERs.

We also demonstrated how a threshold may occur at one ecological scale, but not another. Algal specific growth rate, an organismal physiological response, did not demonstrate a threshold response to nutrient identity or magnitude (Figure 5a-d). In contrast, algal biomass, an ecosystem response, demonstrated a threshold response in three of the four simulations (Figure 5a-d). This discrepancy across scales occurs because algal specific growth rate is dictated by dilution rate in the model (i.e., lake water residence time) rather than stoichiometry. When nutrients become increasingly scarce, individual algae quickly reduce the available nutrients to maintain their growth rate. This maintenance of growth rate causes less biomass to be accrued (i.e., a smaller number of individuals in the population or community). This result highlights the importance of properly identifying the process that is limited. Here, the organismal growth rate is not limited, rather, the ecosystem level process of biomass accrual is limited (Reynolds 1992).

Using the low P simulation, we show that other mechanisms acting across scales can influence the response type, sensitivity, and location of a threshold via their direct or indirect effects on shifts in resource limitation (Figure 6). Parameters acting on the supply of nutrients and light, such as dilution rate and maximum lake depth, did not change the x-value of the stoichiometric threshold, but did change the response type and y-value of the threshold (Figure 6a & b). Parameters which act on algal nutrient demand (algal quota and P half-saturation constant) altered both the response type and x-value of the threshold (Figure 6c & d). Algal physiological traits that dictate nutrient demand were the only parameters which altered the x-value of the threshold. In addition to the identity and magnitude of element(s) that are altering the stoichiometric gradient (Figure 5), care must be taken to identify and explore the supply and demand mechanisms impacting the location of the TER and the shape of its response.

Using well-established theory, the lake supply N:P-algal biomass TER provides a robust example for ecological scaling of the TER which could be expanded upon with other model formulations, organisms, and ecosystems. For example, a model structure with the Droop formulation of algal physiology, which allows for flexible algal stoichiometry, may be warranted to properly capture TER dynamics (Supplementary Information; Droop 1968). Future work could consider alternate model structures and/or designing an empirical experiment to test and compare the model predictions of static vs. flexible algal stoichiometry. A process model approach allows for the development of *a priori* hypotheses regarding the response type and location of a TER that can then be empirically tested. Additionally, such an approach can explore potential mechanisms behind the TER at various scales. A TER is likely driven by some shift in, or release from, energy and nutrient limitation. Process models can aid in our consideration of experimental design when testing for such thresholds. For example, they can help identify keystone elements of systems, and thus, the appropriate stoichiometric gradient required to observe such phenomena. Process models can also help identify the most important drivers involved in modifying the response type and location of the threshold that could then be manipulated or observed in field experiments.

### Considerations for testing and inference from ecological scaling of the TER

We recommend that future studies seeking to broaden the TER consider data requirements, modeling and transformation options, and scale. Making a distinction between the presence vs. absence of a stoichiometric threshold will be critical to scaling the TER. The absence of a threshold may be due to either a non-stoichiometric driver (e.g., light or temperature limitation), or the response is linear, suggesting either the absence of a threshold or that the stoichiometric gradient is not exhaustive enough to determine the threshold. Additionally, a testable TER must be derived from a continuous gradient of a stoichiometric driver as opposed to a binary study (e.g., comparisons above and below some threshold) because gradients are necessary to test for nonlinearity (Kreyling et al. 2014). Such gradients may be difficult to achieve at relatively larger spatial and temporal scales (macroecology and macroevolution), but data synthesis and collaboration across disciplines have the potential to meet this need (Collins et al. 2017). Furthermore, rigorous statistical testing will be necessary to examine whether thresholds exist, including threshold analysis, breakpoint analysis, or formal model fit comparisons (see Dodds et al. 2010; Spake et al. 2022). In addition, logarithmic transformation is recommended for ratio data in ecological stoichiometry (Isles 2020). By linearizing the response curve, logarithmic transformations may visually conceal a threshold, but underlying nonlinearity may still be consistent with predictions. Statistical models such as threshold analysis should be robust to transformation, but this challenge warrants consideration given that the sensitivity to transformation may depend on the specific response type and model structure.

Studies examining TERs at varying scales will need to collect elemental data (which many are doing already; Table 1, Table S1) and distinguish between stoichiometric vs. non-stoichiometric drivers. Alongside mechanistic experiments, explicitly measuring multiple elements will address whether elemental ratios are a signature of the driver or are themselves the driver of the response. Similarly, studies will need to consider whether one element in the ratio, vs. both or even multiple elements, are ultimately responsible for thresholds. Such approaches would ideally be able to examine responses to altered elemental concentrations separately from altered elemental ratios (Kominoski et al. 2015).

There appears to have been greater empirical progress examining ecological compared to evolutionary TERs, highlighting the urgency for more work regarding the influence of stoichiometric change in evolutionary opportunities and innovation (Figure 2, El-Sabaawi et al. 2023). In a number of examples, changing resource stoichiometry appears to be associated with nonlinear shifts in evolutionary rate, but clear evidence supporting this hypothesis is lacking (Table 1, Figure 2). This lack of support is common when attempting to link ecosystem-scale processes to evolutionary changes (Figure 2) and represents an opportunity for future research. In particular, experimental evolution studies paired with observational field evolutionary data could provide evidence to either support or refute the role of TERs in evolutionary processes. Selection of organisms which have relatively short generation times and a strong response to contrasts in resource stoichiometry could yield evidence at the microevolutionary scale. At the macroevolutionary scale, phylogenetic analyses paired with empirical support for stoichiometry as the driving mechanism for speciation could provide evidence to support or refute TERs operating at the macroevolutionary scale.

## Conclusion

Collectively, we show that TERs are pertinent across the ecological hierarchy and evolutionary processes. We also present evidence that such stoichiometric driven thresholds can express counter-intuitive scaling patterns, underscoring the need to better understand mechanisms underpinning TERs and how such phenomena govern non-linearity in ecology and evolution. Given the potential for ecological scaling of TERs, our group has provided recommendations for experimental and analytical approaches to encourage further development of and gain insight on the applicability of TER scaling. Moving forward we recommend that researchers examining stoichiometric driven thresholds be thoughtful about which ecological scales they are examining, scale dependency, and how they are considering the role of elemental ratios in ecological and evolutionary processes.

## Acknowledgements

We thank Sarah Collins, Steve Thomas, and David Nguyen for dialogue. We thank R. Farrell for assistance with literature review.

## Declarations

### Ethics Approval

This article does not contain any studies with human participants or animals performed by any of the authors.

### Consent to participate

For this type of study formal consent is not required.

### Consent for publication

For this type of study formal consent is not required.

### Availability of data and material

all data can be accessed at the published repository: https://doi.org/10.5281/zenodo.10631796.

### Code availability

Full code for the model can be accessed at the published repository: https://doi.org/10.5281/zenodo.10631796.

### Authors’ contributions

JRC, EKM, HMH, ACK, CEW, CRO, BBT, conceived the ideas; CRO, BBT analyzed the data; LAR, UVCO, ENW, BN collected data; BBT, CRO, JRC, EKM, HMH, ACK, CEW, FSO wrote the manuscript; other authors provided editorial advice.

## References

Bergstrom, A.K. (2010) The use of TN:TP and DIN:TP ratios as indicators for phytoplankton nutrient limitation in oligotrophic lakes affected by N deposition. Aquat Sci. 72:277–281

Brasier, M. D., & Callow, R. H. (2007) Changes in the patterns of phosphatic preservation across the Proterozoic-Cambrian transition. Memoirs of the Association of Australasian Palaeontologists. 34: 377–389

Brennan, S. T., Lowenstein T. K., & Horita J. (2004) Seawater chemistry and the advent of biocalcification. Geology 32: 473–476

Bunting, L., Leavitt, P. R., Gibson, C. E., McGee, E. J., & Hall, V. A. (2007) Degradation of water quality in Lough Neagh, Northern Ireland, by diffuse nitrogen flux from a phosphorus-rich catchment. Limnol. Oceanogr. 52: 354–369

Buratti FM, Manganelli M, Vichi S, Stefanelli M, Scardala S, Testai E, Funari E. (2017) Cyanotoxins roducing organisms, occurrence, toxicity, mechanism of action and human health toxicological risk evaluation. Arch Toxicol. 91:1049–1130 doi: 10.1007/s00204-016-1913-6.

Capps, K. A., & Flecker, A. S. (2013) Invasive aquarium fish transform ecosystem nutrient dynamics. Proc. R. Soc. London, B, 280 20131520.

Campbell, I. H., & Squire, R. J. (2010) The mountains that triggered the Late Neoproterozoic increase in oxygen: The Second Great Oxidation Event. Geochimica et Cosmochimica Acta 74: 4187–4206. 10.1016/j.gca.2010.04.064

Cohen, P. A., Strauss, J. V., Rooney, A. D., Sharma, M., & Tosca, N. (2017) Controlled hydroxyapatite biomineralization in an ∼810 million-year-old unicellular eukaryote. Sci Adv. 1700095

Collins, S. M., Oliver, S. K., Lapierre, J. F., Stanley, E. H., Jones, J. R., Wagner, T., & Soranno, P. A. (2017) Lake nutrient stoichiometry is less predictable than nutrient concentrations at regional and sub-continental scales. Ecol App. 27: 1529–1540

Cook, P. J. (1992) Phosphogenesis around the Proterozoic-Phanerozoic transition. J. of the Geol. Soc. 149: 615–620

Delgado-Baquerizo, M., Reich, P. B., Khachane, A. N., Campbell, C. D., Thomas, N., Freitag, T. E., Abu Al-Soud, W., Sorensen, S., Bardgett, R. D., & Singh, B. K. (2017) It is elemental: soil nutrient stoichiometry drives bacterial diversity. Environ. Microbiol, 19: 1176–1188

Devotta, D. A., Fraterrigo, J. M., Walsh, P. B., Lowe, S., Sewell, D. K., Schindler, D. E., & Hu, F. S. (2021) Watershed Alnus cover alters N: P stoichiometry and intensifies P limitation in subarctic streams. Biogeochemistry 153: 155–176

Dodds, W. K., Clements, W. H., Gido, K., Hilderbrand, R. H., & King, R. S. (2010) Thresholds, breakpoints, and nonlinearity in freshwaters as related to management. J N AM Benthol Soc. 29: 988–997

Downing, J.A., & McCauley E. (1992) The nitrogen:phosphorus relationship in lakes. Limnol. Oceanogr. 37:936–945

Droop, M.R. (1968) Vitamin B12 and marine ecology. IV. The kinetics of uptake, growth, and inhibition in Monochrysis lutheri. J. of the Mar. Biol. Assoc. U.K. 48:689–733

El-Sabaawi, R. W., Zandona, E., Kohler, T. J., Marshall, M. C., Moslemi, J. M., Travis, J., López-Sepulcre, A., Rerriére, R., Pringle, C. M., Thomas, S. A., Reznick, D. N., & Flecker, A. S. (2012) Widespread intraspecific organismal stoichiometry among populations of the Trinidadian guppy. Funct. Ecol. 26:666–676

El-Sabaawi, R. W., Lemmen, K. D., Jeyasingh, P. D., & Declerck, S. A. (2023) SEED: A framework for integrating ecological stoichiometry and eco-evolutionary dynamics. Ecol Lett. 26: S109–S126

Elser, J. J., Chrzanowski, T. H., Sterner, R. W., & Mills, K. H. (1998) Stoichiometric constraints on food-web dynamics: a whole-lake experiment on the Canadian Shield. Ecosystems 1: 120–136

Elser, J. J., & Urabe, J. (1999) The stoichiometry of consumer-driven nutrient recycling: theory, observations, and consequences. Ecology 80: 735–751

Elser, J. (2006). Biological stoichiometry: a chemical bridge between ecosystem ecology and evolutionary biology. Amer. Naturalist 168: S25–S35

Elser, J. J., Watts, J., Schampel, J. H., & Farmer, J. (2006) Early Cambrian food webs on a trophic knife-edge? A hypothesis and preliminary data from a modern stromatolite-based ecosystem. Ecol Lett. 9: 295–303

Elser, J. J., Bracken, M. E., Cleland, E. E., Gruner, D. S., Harpole, W. S., Hillebrand, H., Ngai, J.T., Seabloom, E. W., Shurin, J. B., & Smith, J. E. (2007) Global analysis of nitrogen and phosphorus limitation of primary producers in freshwater, marine and terrestrial ecosystems. Ecol Lett. 10: 1135–1142

Enríquez, S. C. M. D., Duarte, C. M., & Sand-Jensen, K. (1993) Patterns in decomposition rates among photosynthetic organisms: the importance of detritus C: N: P content. Oecologia 94: 457–471

Fernàndez-Busquets, 2010. Cambrian Explosion 10.1002/9780470015902.a0022875

Frisch, D., Morton, P. K., Chowdhury, P. R., Culver, B. W., Colbourne, J. K., Weider, L. J., & Jeyasingh, P. D. (2014) A millennial-scale chronicle of evolutionary responses to cultural eutrophication in Daphnia. Ecol Lett. 17: 360–368

Frost, P. C., Benstead, J. P., Cross, W. F., Hillebrand, H., Larson, J. H., Xenopoulos, M. A., & Yoshida, T. (2006). Threshold elemental ratios of carbon and phosphorus in aquatic consumers. Ecol Lett. 9: 774–779

Gibson, C. A., & O’Reilly, C. M. (2012) Organic matter stoichiometry influences nitrogen and phosphorus uptake in a headwater stream. Freshw. Sci. 31: 395–407

Gould, S. J., Pradeu, T., Ricklefs, R. E., Webb, T. J., Gaston, K. J., Rinkevich, B., Forsdyke, D. R., & Rinkevich, B. (1998) Gulliver’s Further Travels: The Necessity and Difficulty of a Hierarchical Theory of Selection. Philos Trans R Soc Lond B Biol Sci 353: 307–314. 56481

Hayes, N. M., Patoine, A., Haig, H. A., Simpson, G. L., Swarbrick, V. J., Wiik, E., & Leavitt, P. R. (2019) Spatial and temporal variation in nitrogen fixation and its importance to phytoplankton in phosphorus-rich lakes. Freshw. Biol. 64: 269–283. 10.1111/fwb.13214

Helfield, J. M., & Naiman, R. J. (2002) Salmon and alder as nitrogen sources to riparian forests in a boreal Alaskan watershed. Oecologia 133: 573–582

Hiatt, D. L., Robbins, C. J., Back, J. A., Kostka, P. K., Doyle, R. D., Walker, C. M., Rains, M. C., Whigham, D. F., & King, R. S. (2017) Catchment-scale alder cover controls nitrogen fixation in boreal headwater streams. Freshw. Sci. 36: 523–532

Isles, P. D. (2020) The misuse of ratios in ecological stoichiometry. Ecology, 101: e03153.

Klausmeier, C.A., Litchman, E., Daufresne, T., & Levin, S.A. (2004) Optimal nitrogen-to-phosphorus stoichiometry of phytoplankton. Nature 429:171–174

Klausmeier, C.A., Litchman, E., Daufresne, T., & Levin, S.A. (2008) Phytoplankton stoichiometry. Ecological Research 23:479–485

Kramer, B. J., Hem, R., & Gobler, C. J. (2022). Elevated CO2 significantly increases N2 fixation, growth rates, and alters microcystin, anatoxin, and saxitoxin cell quotas in strains of the bloom-forming cyanobacteria, Dolichospermum. Harmful Algae, 120, 102354

Kominoski, J. S., Rosemond, A. D., Benstead, J. P., Gulis, V., Maerz, J. C., & Manning, D. W. (2015) Low-to-moderate nitrogen and phosphorus concentrations accelerate microbially driven litter breakdown rates. Ecol Apps, 25:856–865

Kreyling, J., Jentsch, A., & Beier, C. (2014) Beyond realism in climate change experiments: gradient approaches identify thresholds and tipping points. Ecol Lett 17, 125–e1

Laspoumaderes, C., Meunier, C. L., Magnin, A., Berlinghof, J., Elser, J. J., Balseiro, E., Torres, G., Modenutti, B., Tremblay, N., & Boersma, M. (2022) A common temperature dependence of nutritional demands in ectotherms. Ecol Lett 25: 2189–2202

López-Rodas, V., Perdigones, N., Marvá, F., Rouco, M., & García-Cabrera, J. A. (2008) Adaptation of phytoplankton to novel residual materials of water pollution: An experimental model analysing the evolution of an experimental microalgal population under formaldehyde contamination. Bull. Environ. Contam. and Toxicol. 80: 158–162

Manzoni, S., Taylor, P., Richter, A., Porporato, A., & Ågren, G. I. (2012) Environmental and stoichiometric controls on microbial carbon-use efficiency in soils. New Phytologist, 196: 79–91

Martin, M. W., Grazhdankin, D. V., Bowring, S. A., D. Evans, D. A., Fedonkin, M. A., & Kirschvink, J. L. (2000) Age of Neoproterozoic Bilatarian Body and Trace Fossils, White Sea, Russia: Implications for Metazoan Evolution. Science 288: 841–845

Moody, E. K., Lujan, N. K., Roach, K. A., & Winemiller, K. O. (2019). Threshold elemental ratios and the temperature dependence of herbivory in fishes. Funct Ecol. 33, 913–923.

Moody, E. K., Butts, T. J., Fleck, R., Jeyasingh, P. D., & Wilkinson, G. M. (2022) Eutrophication-driven eco-evolutionary dynamics indicated by differences in stoichiometric traits among populations of Daphnia pulicaria. Freshw Bio: 67, 353–364

Mooshammer, M., Wanek, W., Zechmeister-Boltenstern, S., & Richter, A. A. (2014) Stoichiometric imbalances between terrestrial decomposer communities and their resources: mechanisms and implications of microbial adaptations to their resources. Frontiers in microbiology, 5: 22

Osburn, F. S., Wagner, N. D., Taylor, R. B., Chambliss, C. K., Brooks, B. W., & Scott, J. T. (2022) The effects of salinity and N: P on N-rich toxins by both an N-fixing and non-N-fixing cyanobacteria. Limnology and Oceanography Letters, 8: 162–172

Olsen, G. J., Lane, D. J., Giovannoni, S. J., Pace, N. R., & Stahl, D. A. (1986) Microbial ecology and evolution: a ribosomal RNA approach. Annu. Rev. microbiol 40:337–365

Pillon, Y., Hopkins, H. C., Rigault, F., Jaffré, T., & Stacy, E. A. (2014) Cryptic adaptive radiation in tropical forest trees in New Caledonia. New Phytologist, 202:521–530

Plath, K., & Boersma, M. (2001) Mineral limitation of zooplankton: stoichiometric constraints and optimal foraging. Ecology, 82: 1260–1269

Rabosky, D. L., & McCune, A. R. (2010) Reinventing species selection with molecular phylogenies. Trends Ecol Evol. 25: 68–74. 10.1016/j.tree.2009.07.002

Ratajczak, Z., Carpenter, S. R., Ives, A. R., Kucharik, C. J., Ramiadantsoa, T., Stegner, M. A., Williams, J. W., Zhang, J., & Turner, M. G. (2018) Abrupt change in ecological systems: inference and diagnosis. Trends Ecol Evol. 33(7), 513–526

Redfield, A.C. (1958) The biological control of chemical factors in the environment. American Scientist 46:205–221

Reynolds, C. S. (1992) Eutrophication and the management of planktonic algae: what Vollenweider couldn’t tell us. In Sutcliffe, D. W. and Jones, J. G. (eds), Eutrophication: Research and Application to Water Supply. Freshwater Biological Association, Ambleside, pp. 4–29.

Salmon, V. G., Breen, A. L., Kumar, J., Lara, M. J., Thornton, P. E., Wullschleger, S. D., & Iversen, C. M. (2019) Alder distribution and expansion across a tundra hillslope: implications for local N cycling. Frontiers in plant science, 10, 1099

Schade, J. D., MacNeill, Keely., Thomas, S. A., Camille McNeely, F., Welter, J. R., Hood, J., Goodrich, M., Power, M. E., & Finlay, J. C. (2011) The stoichiometry of nitrogen and phosphorus spiralling in heterotrophic and autotrophic streams. Freshw Biol. 3: 424–436.

Shaftel, R. S., King, R. S., & Back, J. A. (2012) Alder cover drives nitrogen availability in Kenai lowland headwater streams, Alaska. Biogeochemistry 107: 135–148

Scheffer, M., & Jeppesen, E. (2007). Regime shifts in shallow lakes. Ecosystems 10, 1–3.

Simons, A. M. (2002) The continuity of microevolution and macroevolution. J. Evol. Biol. 15, 688–701

Smith, M. P., & T. Harper, D. A. (2013) Causes of the Cambrian Explosion. Science 341: 1355–1356

Spake, R., Barajas-Barbosa, M. P., Blowes, S. A., Bowler, D. E., Callaghan, C. T., Garbowski, M., Jurburg, S. D., van Klink, R., Korell, L., Ladouceur, E., Rozzi, R., Viana, D.S., Xu, W., & Chase, J. M. (2022) Detecting thresholds of ecological change in the Anthropocene. Annu. Rev. of Environ. Resour. 47: 797–821.

Squire, R. J., Campbell, I. H., Allen, C. M., & Wilson, C. J. (2006) Did the Transgondwanan Supermountain trigger the explosive radiation of animals on Earth? Earth Planet. Sci. Lett. 250:116–133. 10.1016/j.epsl.2006.07.032

Stelzer, R. S., & Lamberti, G. A. (2001) Effects of N: P ratio and total nutrient concentration on stream periphyton community structure, biomass, and elemental composition. Limnol. Oceanogr. 46: 356–367

Sterner, R.W. & Elser, J.J. (2002) Ecological stoichiometry: the biology of elements from molecules to the biosphere. Princeton, New Jersey: Princeton University press.

Stottlemyer, R., & Toczydlowski, D. (1999) Seasonal relationships between precipitation, forest floor, and streamwater nitrogen, Isle Royale, Michigan. Soil Sci Soc Am J. 63: 389–398

Su, H., Wu, Y., Xia, W., Yang, L., Chen, J., Han, W., Fang, J., & Xie, P. (2019) Stoichiometric mechanisms of regime shifts in freshwater ecosystem. Water research, 149: 302–310

Taylor, P. G., & Townsend, A. R. (2010) Stoichiometric control of organic carbon–nitrate relationships from soils to the sea. Nature 464: 1178–1181

Tilman D (ed). 1982. Resource competition and community structure. Princeton University Press, Princeton, NJ.

Tromboni, F., Thomas, S. A., Gücker, B., Neres-Lima, V., Lourenço-Amorim, C., Moulton, T. P., Silva-Junior, E. F., Reijó-Lima, R., Boëchat, I. G., & Zandonà, E. (2018) Nutrient limitation and the stoichiometry of nutrient uptake in a tropical rain forest stream. Journal of Geophysical Research: Biogeosciences 123, 2154–2167.

Urabe, J., & Watanabe, Y. (1992) Possibility of N or P limitation for planktonic cladocerans: an experimental test. Limnol. Oceanogr. 37, 244–251

Van de Waal, D. B., Verspagen, J. M., Lürling, M., Van Donk, E., Visser, P. M., & Huisman, J. (2009) The ecological stoichiometry of toxins produced by harmful cyanobacteria: an experimental test of the carbon-nutrient balance hypothesis. Ecol lett. 12, 1326–1335.

Van de Waal, D. B., Smith, V. H., Declerck, S. A., Stam, E. C., & Elser, J. J. (2014) Stoichiometric regulation of phytoplankton toxins. Ecol lett. 17: 736–742

Vitousek, P. M., Cassman, K. E. N., Cleveland, C., Crews, T., Field, C. B., Grimm, N. B., Howarth, R. W., Marino, R., Martinelli, L., Rastetter, E. B., & Sprent, J. I. (2002) Towards an ecological understanding of biological nitrogen fixation. The nitrogen cycle at regional to global scales, 1-45

Volk, C. J., Kiffney, P. M., & Edmonds, R. L. (2008) Nutrient limitation in red alder (Alnus rubra) and conifer forested streams of western Washington state, USA. Amer Midl Naturalist 159: 190–199

Wagner, N. D., Osburn, F. S., Taylor, R. B., Back, J. A., Chambliss, C. K., Brooks, B. W., & Scott, J. T. (2023) Diazotrophy modulates cyanobacteria stoichiometry through functional traits that determine bloom magnitude and toxin production. Limnol. Oceanogr. 68: 348–360.

Yan, Z., Han, W., Peñuelas, J., Sardans, J., Elser, J. J., Du, E., Reich, P. B., & Fang, J. (2016) Phosphorus accumulates faster than nitrogen globally in freshwater ecosystems under anthropogenic impacts. Ecol lett. 19: 1237–1246. 10.1111/ele.12658

Zhou, J., Han, X., Brookes, J. D., & Qin, B. (2022) High probability of nitrogen and phosphorus co-limitation occurring in eutrophic lakes. Environmental Pollution, 292: 118276. 10.1016/j.envpol.2021.118276

